# Cell-type-specific effects of autism-associated chromosome 15q11.2-13.1 duplications in human brain

**DOI:** 10.1101/2024.05.22.595175

**Authors:** Caroline Dias, Alisa Mo, Chunhui Cai, Liang Sun, Kristen Cabral, Catherine A. Brownstein, Shira Rockowitz, Christopher A. Walsh

**Affiliations:** Department of Pediatrics, Section of Developmental Pediatrics, Section of Genetics and Metabolism, Children’s Hospital Colorado, University of Colorado Anschutz Medical Campus, Aurora, CO 80045; Division of Developmental Medicine, Boston Children’s Hospital, Boston, MA 02115; Division of Genetics and Genomics, Manton Center for Orphan Disease Research, Boston Children’s Hospital, Boston, MA 02115; Department of Pediatrics, Harvard Medical School, Boston, MA 02115; Department of Neurology, Boston Children’s Hospital, Harvard Medical School, Boston, MA 02115; Research Computing, Department of Information Technology, Boston Children’s Hospital, Boston, MA 02115; Howard Hughes Medical Institute, Boston Children’s Hospital, Boston, MA 02115

**Keywords:** dup15q, autism, snRNA-seq, ATAC-seq, copy number variant, neurodevelopment

## Abstract

Recurrent copy number variation represents one of the most well-established genetic drivers in neurodevelopmental disorders, including autism spectrum disorder (ASD). Duplication of 15q11.2-13.1 (dup15q) is a well-described neurodevelopmental syndrome that increases the risk of ASD by over 40-fold. However, the effects of this duplication on gene expression and chromatin accessibility in specific cell types in the human brain remain unknown. To identify the cell-type-specific transcriptional and epigenetic effects of dup15q in the human frontal cortex we conducted single-nucleus RNA-sequencing and multi-omic sequencing on dup15q cases (n=6) as well as non-dup15q ASD (n=7) and neurotypical controls (n=7). Cell-type-specific differential expression analysis identified significantly regulated genes, critical biological pathways, and differentially accessible genomic regions. Although there was overall increased gene expression across the duplicated genomic region, cellular identity represented an important factor mediating gene expression changes. Neuronal subtypes, showed greater upregulation of gene expression across a critical region within the duplication as compared to other cell types. Genes within the duplicated region that had high baseline expression in control individuals showed only modest changes in dup15q, regardless of cell type. Of note, dup15q and ASD had largely distinct signatures of chromatin accessibility, but shared the majority of transcriptional regulatory motifs, suggesting convergent biological pathways. However, the transcriptional binding factor motifs implicated in each condition implicated distinct biological mechanisms; neuronal JUN/FOS networks in ASD vs. an inflammatory transcriptional network in dup15q microglia. This work provides a cell-type-specific analysis of how dup15q changes gene expression and chromatin accessibility in the human brain and finds evidence of marked cell-type-specific effects of this genetic driver. These findings have implications for guiding therapeutic development in dup15q syndrome, as well as understanding the functional effects CNVs more broadly in neurodevelopmental disorders.

## Introduction

The proximal end of the long arm of chromosome 15 is a genomic region containing several segmental duplications, making it particularly susceptible to complex rearrangements at recurrent breakpoints (Fig 1A) ^1^. Parental chromosome-specific deletions in this region lead to the imprinting disorders Angelman syndrome and Prader-Willi syndrome, whereas maternal duplication of 15q11-13, referred to hereafter as dup15q syndrome, is associated with autism spectrum disorder (ASD), intellectual disability, hypotonia and epilepsy ^2^. There is over a 40-fold increased risk of ASD in individuals carrying this duplication, making it one of the most significant and highly penetrant genetic drivers of ASD ^3,4^.

**Figure 1:**
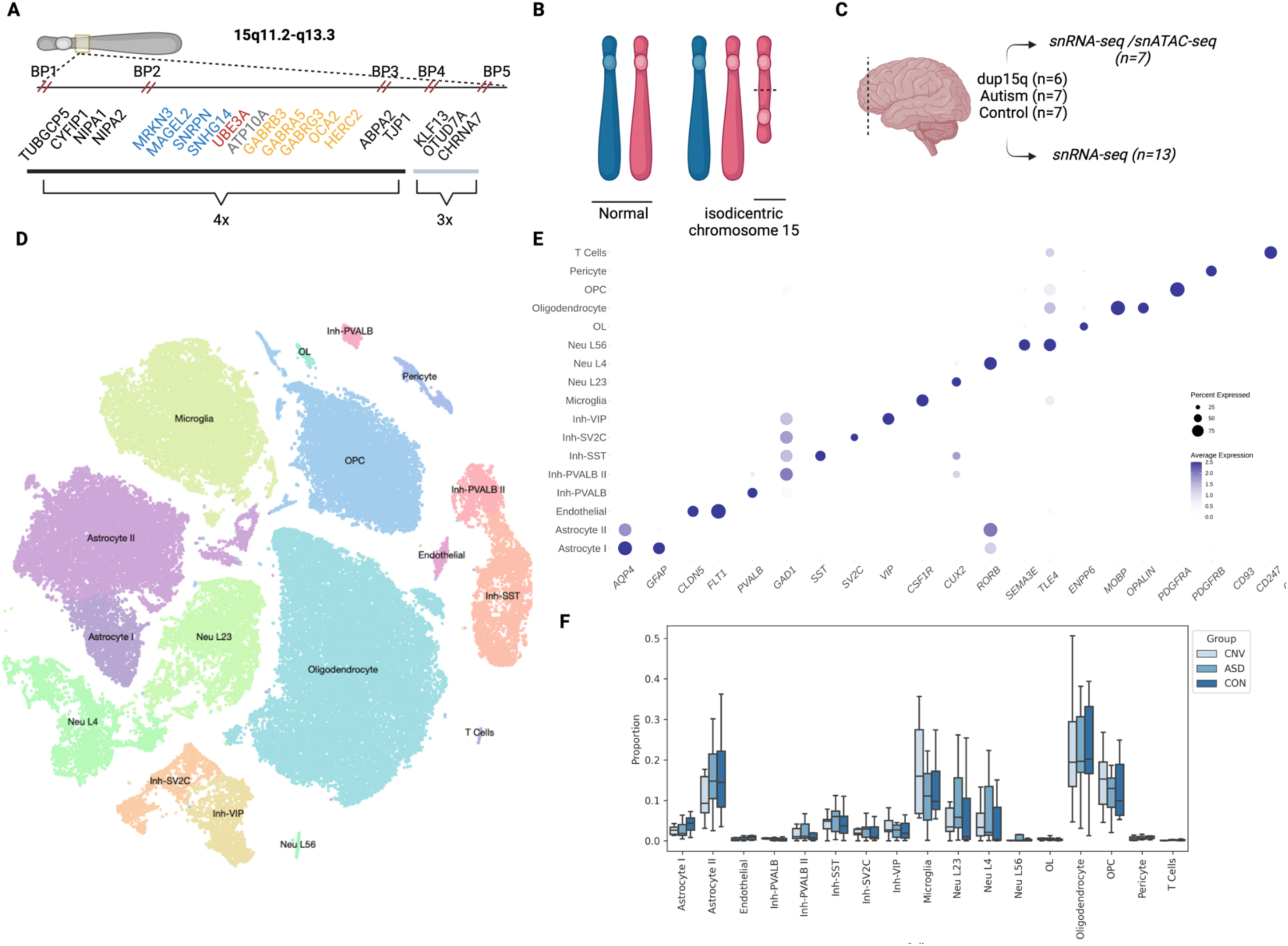
Single nuclei analysis of dup15q in human brain **A.** Genomic architecture of chromosome 15 with only select genes shown for clarity. Inset shows location of genes with breakpoints and copy number indicated. Critical imprinting locus between BP 2 and 3-Blue: paternally imprinted, red: maternally imprinted, yellow:bi-allelic, grey: conflicting reports. **B.** Cartoon representation of dup15q structure due to isodicentric maternal chromosome 15. **C.** Samples used from frontal cortex. **D.** Visualization of single-nuclei data with t-SNE dimension reduction. **E.** Cell-type-specific gene expression indicate representation of expected cell types in human frontal cortex and accurate clustering. **F.** Cell-type compositional analysis indicates no significant changes in proportion of cell types between conditions. Boxplot shows median, interquartile range (box), and +/- 1.5 interquartile range (error bars) of proportion of each cell type.

Most cases of dup15q are caused by an isodicentric chromosome 15 (Fig 1B)^5^, and many of these are in turn driven by an asymmetric recombination event between breakpoints 4 and 5. This leads to tetrasomy from the centromere to breakpoint 4 (a region that includes the Prader-Willi/Angelman critical region [PWACR] between breakpoints 2 and 3) and trisomy from breakpoint 4 to 5 (Fig 1A) ^2^. Given that most cases are caused by maternally-derived duplications and that the duplication contains a critical imprinting locus within the PWACR, copy number alterations in maternally imprinted, dosage sensitive genes such as *UBE3A* have been hypothesized to underly the dup15q clinical phenotype. However, the dup15q region also contains a GABA receptor cluster and some individuals with dup15q have atypical responses to benzodiazepines, potentially implicating these and other genes as well.

The cell-type-specific effects of dup15q on gene expression in the human brain are unknown. Prior work in post-mortem brain has shown that in bulk tissue, *UBE3A* expression does not match changes in gene dosage in dup15q cases ^6^. However, bulk brain tissue analysis cannot identify whether only specific cell types showed changes in gene expression. Understanding how cell-type-specific heterogeneity in gene expression and chromatin accessibility contributes to pathophysiology is the first step to identify potential therapeutic targets and advance precision medicine for this disorder.

It is also critical to understand the similarities and differences on the molecular and cellular level between dup15q and ASD not related to dup15q. Although prior studies have demonstrated similarities on the molecular level in the brain between individuals with dup15q and ASD, suggesting convergent biological processes^7–9^, there are also important clinical distinctions in the developmental and behavioral profile of these subgroups. Individuals with dup15q syndrome may have an initially preserved social smile and yet more marked neurological co-morbidities, including greater motor impairment ^10,11^. The complexity of this genomic region, in addition with the long-standing challenge of studying the heterogeneity of brain tissue, has made it difficult to directly parse the effects of this variant in human brain. In this study we sought to examine the transcriptional and epigenetic landscape of dup15q brain cell types compared to both neurotypical controls and non-dup15q ASD cases, hypothesizing that cell-type-specific effects, previously masked by brain tissue heterogeneity, would reveal important biological distinctions in cellular pathophysiology and provide foundational information for future mechanistic and translational studies.

## Materials & Methods

### Brain Tissue

Tissue was obtained from the NIH NeuroBioBank and Autism BrainNet according to their institutional review board approvals and following written informed consent. Initial dissection of tissue for brain bank specimens was done under standardized procedures using sequential sectioning. Research on these deidentified specimens and data was performed at Boston Children’s Hospital with approval from the Committee on Clinical Investigation. Post-mortem frozen frontal cortex was obtained from 6 cases with dup15q, 7 non-dup15q ASD cases and 7 neurotypical controls. 13 samples were processed for snRNA-seq and 7 for multi-omic sequencing (Fig 1C). All of the dup15q cases were validated through copy number analysis of the PWACR (tetrasomy) ^6,7^ and/or optical genome mapping (Fig S1). Methylation status in brain has also been previously assessed in the majority of dup15q cases^6,12^. Groups were age and sex-matched (14-17% female) and there were no significant differences between RNA integrity number (RIN) and post-mortem interval (PMI) between groups (Table 1, Fig S2). Frozen tissue was stored at -80 and kept frozen until processing. ∼25 mg pieces of tissue were dissected off the larger tissue block on a pre-chilled cryostat maintained at -20 degrees C, using pre-chilled sterile forceps and scalpels.

**Table 1:** Demographic information for samples. PMI-post-mortem interval. RIN-RNA integrity number.

| ID | Condition | Age | Sex | PMI | RIN | Seizure History |
| --- | --- | --- | --- | --- | --- | --- |
| 797 | ASD | 9 | M | 13 | 7.7 |  |
| 1793 | control | 11 | M | 13 |  |  |
| 1793 | control | 11 | M | 19 |  |  |
| 4643 | control | 42 | F | 4 | 8.2 |  |
| 5023 | ASD | 45 | M |  |  |  |
| 5278 | ASD | 15 | F | 13 | 6.9 | Yes |
| 5391 | control | 8 | M | 19 | 6.9 |  |
| 5408 | control | 6 | M | 12 |  |  |
| 5497 | control | 68 | M | 24 | 8 |  |
| 7TJ5 | ASD | 13 | M | 26 | 6.7 |  |
| 862K | dup15q | 12 | M | 30 | 5.4 | Yes |
| AN05983 | dup15q | 24 | M | 36 | 7.6 | Yes |
| AN06365 | dup15q | 10 | M | 18 | 8.1 | Yes |
| AN09412 | ASD | 29 | M | 38 | 7.3 |  |
| AN11931 | dup15q | 39 | F | 33 | 8.5 | Yes |
| AN14067 | ASD | 38 | M | 46 | 8.7 | Yes |
| AN14762 | dup15q | 9 | M | 14 |  | Yes |
| PGEQ | control | 16 | M | 23 | 6.8 |  |
| 5565 | ASD | 12 | M | 22 |  |  |
| Z2WU | dup15q | 13 | M | 17 | 7.4 | Yes |

### snRNA-sequencing & multi-omic sequencing

For nuclear isolation, tissue was subjected to glass dounce homogenization followed by sucrose gradient centrifugation to isolate nuclei, as previously described ^13^. For snRNA-seq, nuclei were resuspended in a nuclear resuspension buffer (1% BSA in PBS with 1 U/uL RNase inhibitor) and stained with Hoechst immediately prior to fluorescent-activated nuclear sorting on a BD Aria II, during which gating was used to remove debris, dying nuclei, and doublets. 10,000 nuclei/ sample were sorted with a 100 um low pressure nozzle and immediately processed for 10x encapsulation as described below to generate single nuclei libraries. Following encapsulation on the 10X Chromium controller, libraries were prepared as per the 10X Genomics Single Cell 3’ Kit. cDNA and library quality control were ensured by assessing DNA on the Agilent Bioanalyzer High Sensitivity chip.

For generation of multiome libraries, 50,000 nuclei were sorted as above directly into buffer (10mM Tris-HCl pH 7.4, 10 mM NaCl, 3 mM MgCl2, 1% BSA, 1 mM DTT, RNase inhibitor 1U/uL). 1X Lysis Buffer (10 mM Tris-HCl, 10 mM NaCl, 3 mM MgCl2, 0.1% Tween-20, 0.1% NP-40, 0.01% Digitonin, 1% BSA, 1 mM DTT,1 U/uL RNase inhibitor) was added to a final concentration of 0.1X and the nuclei were incubated on ice for 2 minutes. Tween-20 was added (final concentration 0.1%), and nuclei were immediately pelleted by centrifugation at 500g x 5 minutes at 4C in a bucket centrifuge. The pellet was resuspended in Diluted Nuclei Buffer (1X Nuclei Buffer (10X Genomics), 1mM DTT, 1 U/uL RNase inhibitor) and centrifuged. The number of nuclei in the pellet was quantified using a hemocytometer before proceeding with the 10X Genomics Chromium Next GEM Single Cell Multiome ATAC + Gene Expression kit according to manufacturer’s instructions.

Libraries were sequenced with paired-end 150-bp reads on an Illumina NovaSeq 6000.

### Bioinformatics Analysis

For snRNA-seq analysis, the following steps were taken to process the samples: demultiplexing of raw data, alignment (to GRCh382020-A), quality control and filtering (see below), dimensionality reduction and unsupervised clustering. R (v4.2.3), Cell Ranger (v6.1.1.) and Seurat (v4.3.0) ^14^ were used on the Boston Children’s computing cluster E2. SoupX ^15^ was applied to remove ambient RNAs and scds^16^ was used to filter out doublets and cells with extreme library sizes (out of the 95% confidence interval), number of features and high content of mitochondrial reads (>10%). Quality control was performed over each independent sample and ∼ 50-80% nuclei remained following bioinformatic filtering, depending on the sample, with equivalent metrics between conditions (Fig S3). Post-hoc analysis of known cell-type-specific and layer-specific markers were applied to the remaining nuclei to identify clusters and optimize resolution. One cluster representing a small percentage of nuclei (<1%) was manually removed given signatures of ambient RNA contamination as previously reported^17^. There is increased variability in single-nucleus RNA studies from human tissue. Thus, in addition to ensuring no significant differences between age, percent male, RIN, and PMI (Table 1, Fig S2) we also assessed the impact of various demographic factors on gene expression variability. We found minimal impact of age and sex using principal component analysis (Fig S2) and thus conducted differential expression analysis with the FindMarkers functionality in Seurat using default settings of Wilcoxon rank sum test, minimum log fold change > 0.2, padj <.01 for all cell clusters. Our power calculations suggest approximately 400 nuclei/condition are required to detect 80% of gene expression changes with a false discovery rate of 5%. We omitted down-sampling to preserve power. For gene ontology (GO) enrichment, we used clusterProfiler R package ^18,19^. A pathway is treated as enriched if an adjusted p-values (with Benjamin-Hochberg correction) was smaller than 0.05. To identify changes in cellular composition, we used scCODA (single-cell compositional data analysis), a Bayesian hierarchical model developed for analyzing compositional data from single-cell RNA sequencing studies ^20^. It identifies cell types that are differentially abundant between conditions while accounting for the compositional nature of single-cell data.

We employed inferCNV (v1.14.2)^21^, a computational tool designed to infer copy number variation from single-cell RNA sequencing data, to investigate genomic instability across various conditions and cell types using default parameters. We also utilized the R packages NGCHM (v0.13.0) and infercnvNGCHM (v0.1.1) for generating NGCHM files, thereby enhance our inferCNV data interpretation and visualization with the NGCHM (Next-Generation Clustered Heat Map) interactive viewer ^22^.

For multiome analysis, each sample was mapped to the human reference genome (CRCh38-2020-A-2.0.0) using CellRanger Arc (v.2.0.0) with stringent snATAC cell filtering criteria including counts per cell ranging from 1,000 to 1,000,000, nucleosome signal less than 2, and transcription start site (TSS) enrichment score larger than 1. Data preprocessing and normalization were conducted using Signac (v.1.9.0)^23^, while sample integration was achieved through Harmony (v.0.1.1)^24^. Cell type annotations were informed by snRNAseq analysis results. We conducted motif enrichment analysis using Signac, adding motif information to the Seurat object via the AddMotifs function, and identified overrepresented motifs between various conditions using FindMarkers. Motif activities per cell were computed using the chromVAR (v.1.5.0)^25^. For visualization, we generated multiple genomic plot types through Signac, including accessibility tracks and gene annotations, leveraging the CoveragePlot() function for comprehensive genome browser-style presentations. This integrative approach allowed for a nuanced exploration of genomic landscapes, highlighting differential accessibility and gene expression in various cell types.

To generate a list of high-interest genes related to the duplicated region used for visualization and to focus our analysis, we utilized GRCh38/hg38 assembly accessed through genome.ucsc.edu in conjunction with prior publications^1,6,7,26^; given individual differences in precise breakpoints we also included genes beyond breakpoint 5 for comparative purposes.

### Optical Mapping

A custom protocol for human brain tissue was utilized (Bionano Genomics DN30400, Rev A). Briefly, 15-20 mg of brain tissue was homogenized for 10 seconds in Bionano Genomics Homogenization Buffer (P/N 20406) using a TissueRuptor, passed through a 40µm filter, then pelleted at 2000 x g for 5 minutes. The pellet was resuspended and re-pelleted from Bionano Genomics Wash Buffer A (P/N 20407). Homogenized tissue was then digested in 50µL of Protease solution and Bionano Genomics Lysis and Binding Buffer (LBB) (P/N 20375) on a HulaMixer for 15 minutes at 10 rpm. Tissue was further digested in Proteinase K on a HulaMixer for 15 minutes at 10 rpm. Enzymes were inhibited by PMSF.

gDNA binding was performed in the presence of Bionano Genomics Salting Buffer (P/N 20404) and 100% Isopropanol to a Bionano Genomics Nanobind Disk (P/N 20402) on a HulaMixer for 30 minutes at 10 rpm. Nanobinds were washed with Bionano Genomics Wash Buffer 1 (P/N 20376) and Wash Buffer 2 (P/N 20377) (2 rounds each) using a Dynamag Tube Rack. gDNA was eluted from Nanobind using Bionano Genomics Elution Buffer (P/N 20378). gDNA was homogenized for 1 hour at 15 rpm and incubated at room temperature for at least 3 days before labeling and staining. gDNA was fluorescently labeled and stained using the Bionano Genomics Direct Label and Stain Kit (P/N 80005) following Bionano Prep Direct Label and Stain protocol (P/N 30206).

Data was collected on the Saphyr instrument to a target throughput of 1500 Gbp or 400X effective coverage of the GRCh38 reference. Data was submitted to the Rare Variant Analysis pipeline on Bionano Solve version 3.6.1. The Rare Variant Pipeline aligns molecules to the reference to detect relative differences in the sample, forms consensus maps of molecule support in these loci and in the presence of sufficient molecule support, a structural variant call is made. The molecule data was subsequently down-sampled to 400Gbp or 100X coverage of the GRCh38 reference and submitted to the De Novo Assembly pipeline in Bionano Solve version 3.6.1 during which molecules were assembled de novo to form haplotype consensus maps, which were then aligned to the GRCh38 reference for structural variant detection and calling. In both scenarios, global reference coverage is used to determine copy number aberrations in addition to the aforementioned structural variant calling. Results were reviewed in Bionano Access 1.6.1. During review, structural variant and copy number calls were filtered against the Bionano provided mask files which remove structural variant and copy number calls in complex or poorly constructed regions of the GRCh38 reference. Variant calls were filtered against the Bionano control database of healthy individuals retaining only variants present in less than 1% of the control database. Variants were also filtered according to recommended confidence score filtering; 0 for indels, 0.7 for inversions, 0.3 for intrachromosomal fusions, 0.65 for intrachromosomal translocations, and no confidence scores were calculated for duplications where a placeholder value of -1 is used. Variants remaining after filtering were further curated by careful review of the data.

## Results

Following bioinformatic filtering, ∼78,815 high-quality nuclei remained for the combined analysis (Fig 1D). Each condition had comparable numbers of nuclei (Table 2). Post-hoc annotation of cluster types from unsupervised clustering confirmed the expected distribution of layer-specific excitatory neurons, inhibitory neuron subtypes, glia, and other non-neuronal subtypes, that have been previously described in cortical human tissue (Fig 1E) ^13,27–29^. No statistically significant changes in the composition of cell types between conditions was observed (Table 2, Fig 1F), consistent with independent reports ^30^.

**Table 2:** Nuclei number per condition and total nuclei. Green indicates cell type subsequently grouped in “glia” category, and red highlights “neuron”, for ‘low-resolution’ analysis.

| Cell Type | CNV | ASD | CON | Total |
| --- | --- | --- | --- | --- |
| Astrocyte I | 624 | 627 | 1257 | 2508 |
| Astrocyte II | 3214 | 3973 | 4461 | 11648 |
| Endothelial | 107 | 139 | 239 | 485 |
| Inh-PVALB | 158 | 103 | 113 | 374 |
| Inh-PVALB II | 431 | 416 | 473 | 1320 |
| Inh-SST | 1034 | 969 | 1324 | 3327 |
| Inh-SV2C | 699 | 454 | 683 | 1836 |
| Inh-VIP | 871 | 475 | 776 | 2122 |
| Microglia | 4228 | 2929 | 3806 | 10963 |
| Neu L23 | 1869 | 1722 | 2623 | 6214 |
| Neu L4 | 1240 | 1250 | 1733 | 4223 |
| Neu L56 | 32 | 149 | 70 | 251 |
| OL | 110 | 120 | 84 | 314 |
| Oligodendro | 5409 | 5869 | 10888 | 22166 |
| OPC | 3662 | 2916 | 3650 | 10228 |
| Pericyte | 165 | 196 | 313 | 674 |
| T Cells | 33 | 51 | 78 | 162 |
| **Total** | **23886** | **22358** | **32571** | **78815** |

Using all nuclei in a pseudo-bulk tissue analysis demonstrated the expected up-regulation of gene expression in dup15q compared to control within the duplicated region, including the *UBE3A* gene, consistent with prior published bulk RNA-seq work ^7^ (Fig 2A, Supplemental File 1, Supplemental Table 1). We also identified more DEGs globally in dup15q vs control, as compared to ASD vs control or dup15q vs ASD, suggesting unique biological perturbations in dup15q (Supplemental Table 1).

**Figure 2:**
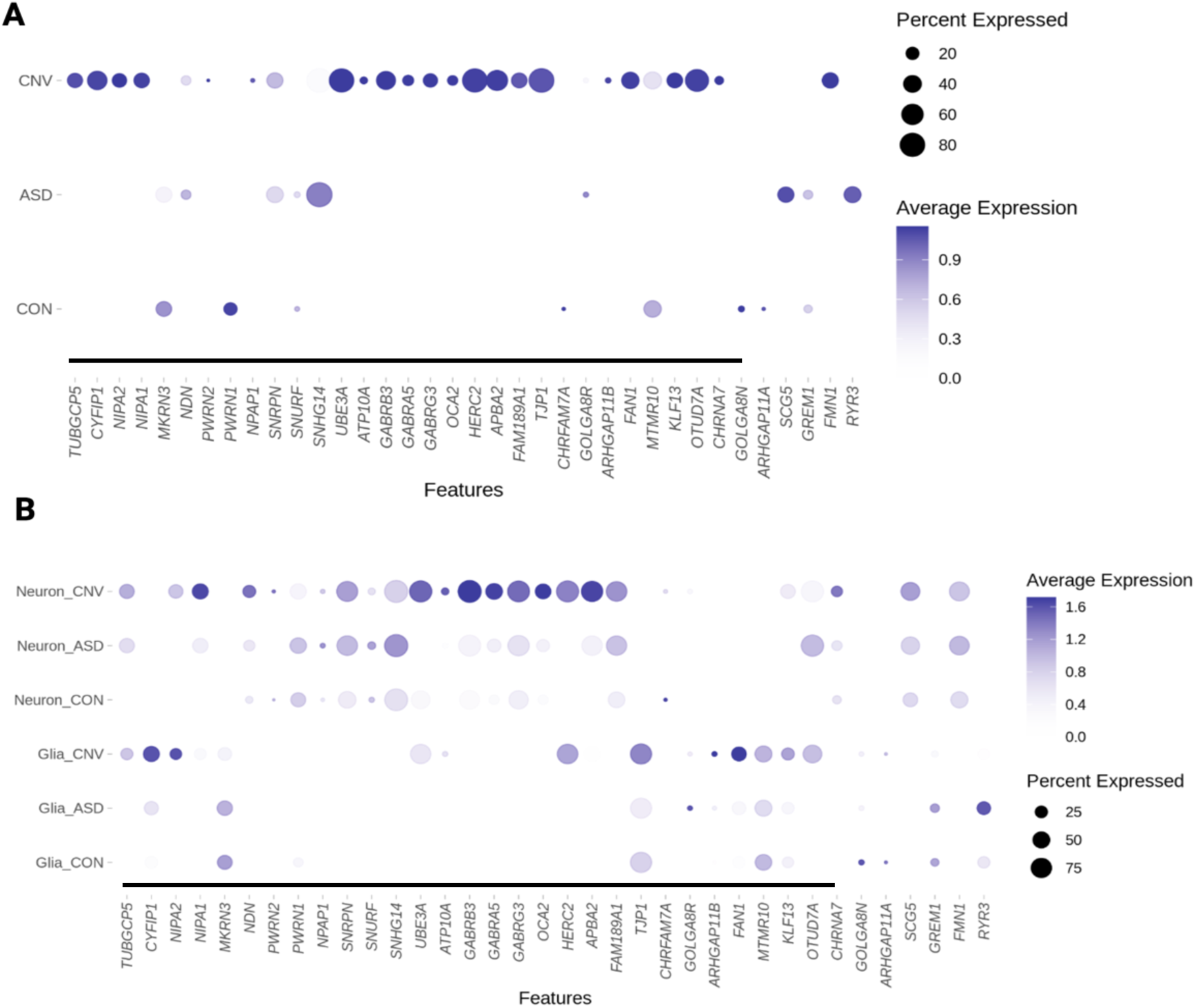
Expression changes within proximal region of chromosome 15q in ASD, 15q (CNV) and control (CON) cases. **A.** Expected upregulation of the duplicated region when examining all nuclei. **B.** Neuronal and glial nuclei broken down by condition demonstrate distinct patterns of changes in gene expression within this region; neurons demonstrate marked upregulation of the PWACR. Color heat map is scaled average expression; black line on bottom demarcates duplication. Note that because visualization includes all cell types using scaled average expression, basal and ceiling effects may mask visualization of some changes between conditions. The list of significant differentially regulated genes can be found in Supplemental Files 4-6.

We then conducted both high and low-resolution differential expression analyses. First, for a ‘low-resolution’ approach, we grouped broad neuronal subtypes (inhibitory and excitatory) and glial subtypes (oligodendrocyte lineage, microglia, and astrocytes). This approach maximized our power to detect expression changes within these categories^31^. Genome-wide, dup15q vs control comparisons demonstrated the largest number of differentially expressed genes, in both neurons and glia, consistent with the broad analysis described above (Supplemental Table 2). Neurons showed more marked increased expression across the duplicated region compared to glia (Fig 2B). Genes within the PWACR region demonstrated preferential upregulation in neurons vs glia in dup15q cases. (Supplemental Files 2-3, Fig 2B). Not all genes demonstrated upregulation in both neurons and glia, although some, like *UBE3A* and *HERC2*, for example, did.

We also conducted “high-resolution” analysis that assessed changes in gene expression in more finely resolved individual cell types (i.e., L2/3 excitatory neurons, microglia, etc.). This analysis reinforced our findings above-namely that cellular context is a critical parameter in gene expression changes, with cell-type-specific effects observed even within neuron or glial subclasses (Supplemental File 4-6, Fig 3). For example, the cluster of GABA receptor genes located within the PWACR demonstrated heterogeneity in gene expression changes between individual neuronal subtypes (Fig 3, Supplemental File 4-6). GABRB3 was upregulated in OPCs and all inhibitory and excitatory neuron subtypes except for layer 5-6 neurons. GABRA5 on the other hand was upregulated only in SV2C inhibitory neurons and layer 2-4 excitatory neurons. Thus, there was unexpected heterogeneity even across similar genes and cell types (Fig 3).

**Figure 3:**
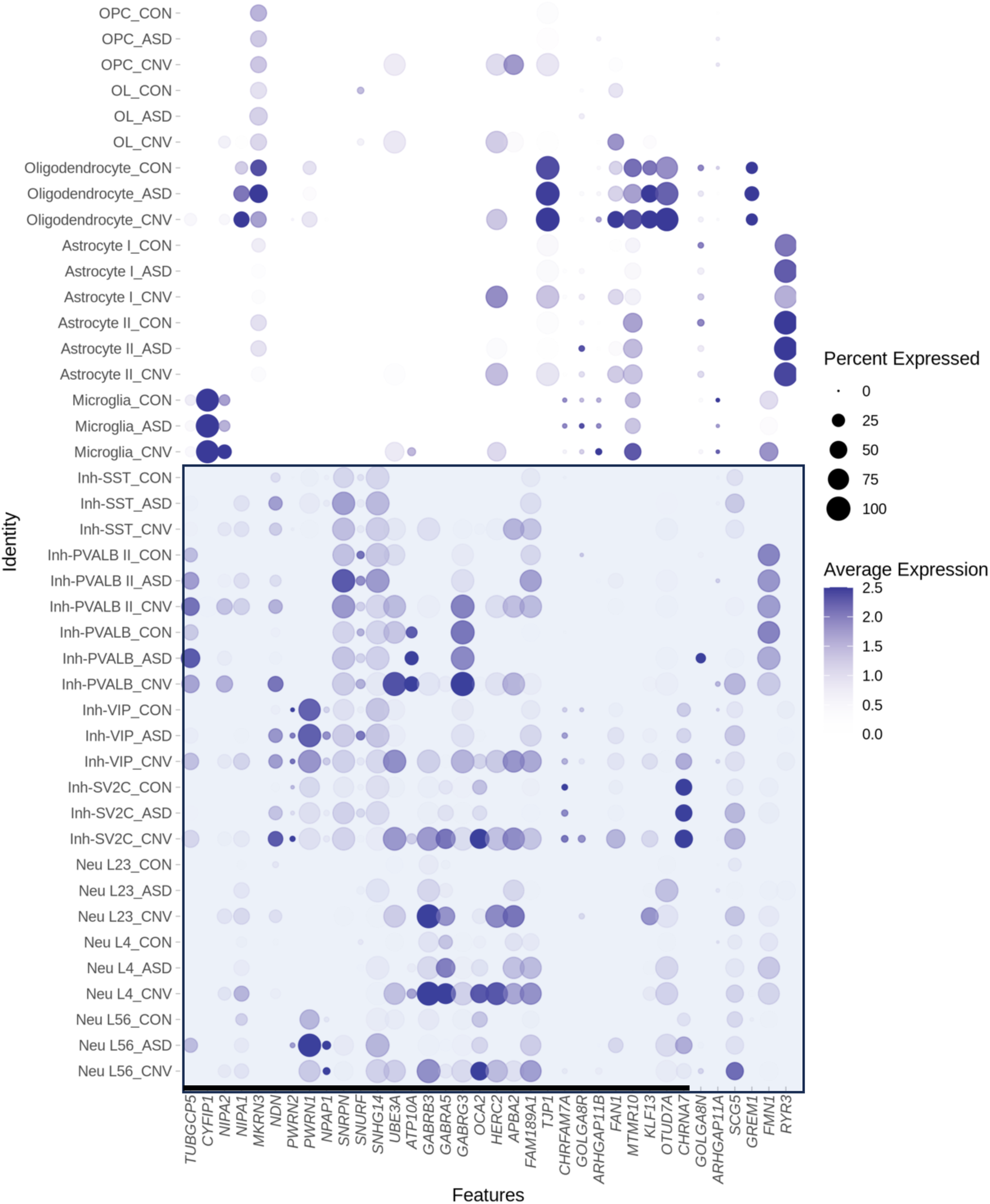
Dot plot depicting global cell-type-specific expression within proximal region of chromosome 15q. Blue box highlights neuronal (bottom) subtypes. Note that because visualization includes all cell types, basal and ceiling effects may mask visualization of some changes between conditions. The list of significant differentially regulated genes can be found in Supplemental Files 4-6.

We took an orthogonal bioinformatic approach to confirm these cell-type-specific changes in gene expression in dup15q cases across the duplication. Bioinformatic tools, such as inferCNV, have been developed to infer somatic copy number variant status from single cell transcriptomic studies in the cancer biology field, in which structural genomic alterations can confer cellular survival advantages and underlie clonal expansion ^21^. We reasoned that although dup15q is a germline event, applying such tools to our data would reveal underlying heterogeneity in gene expression across the duplication. InferCNV applies a hidden Markov model to predict 6 states of gene dosage ranging from complete loss (State 1) to three or more copies (State 6) based on single-cell RNA-sequencing. As expected, applying inferCNV to all dup15q nuclei predicted a higher proportion of high gene dosage states in dup15q as compared to ASD (chi-square p<.001). However, as expected, upregulation across the region of duplication was heterogenous (Fig S5A). This again suggests that there is marked individual variability in dup15q changes and furthermore that not all cell types demonstrate ‘expected’ changes within the duplicated region in dup15q in human brain.

We next sought to identify potential mediating factors of gene expression changes. Differential gene expression studies typically focus on cells in which genes of interest are highly expressed at baseline, given the logic that those would be the cell types critical to understanding cellular pathophysiology; but copy number gains create a situation where cell types with normally low expression may demonstrate greater fold changes in gene expression. We observed such a pattern with *CYFIP1*, located within the duplicated region. *CYFIP1* is normally highly expressed in microglia, and prior work has examined the role of microglial CYFIP1 in neurodevelopment ^32,33^. In all nuclei taken together, *CYFIP1* mRNA was identified as significantly upregulated in dup15q (Supplemental File 1); this upregulation in expression was associated with increases in chromatin accessibility (Fig S4). However, these global changes were not observed in microglia, likely due to its high basal expression (Fig S4). Rather, cell types that demonstrated significant upregulation of *CYFIP1* were all neuronal, including inhibitory VIP and PVALB I and II neurons, and excitatory neurons in layers 2-4 (Supplemental Files 4-6). Thus, our approach reveals that genes previously implicated in neurodevelopment can become mis-expressed in non-canonical cell types in dup15q.

To determine whether baseline expression could be mediating cell-type-specific effects on gene expression within the duplicated region, we examined the relationship between baseline expression in control nuclei and the change in expression in dup15q cases. There was no shift in distribution in the baseline expression of genes of interest in the duplicated region in control nuclei between neurons and glia. Genes with highest baseline expression demonstrated modest change in dup15q cases, a finding robust amongst different cell types. (Fig 4A) Thus, the neuronal specific signature of increased expression was not due to neurons having different baseline gene expression of the genes within this region.

**Figure 4.**
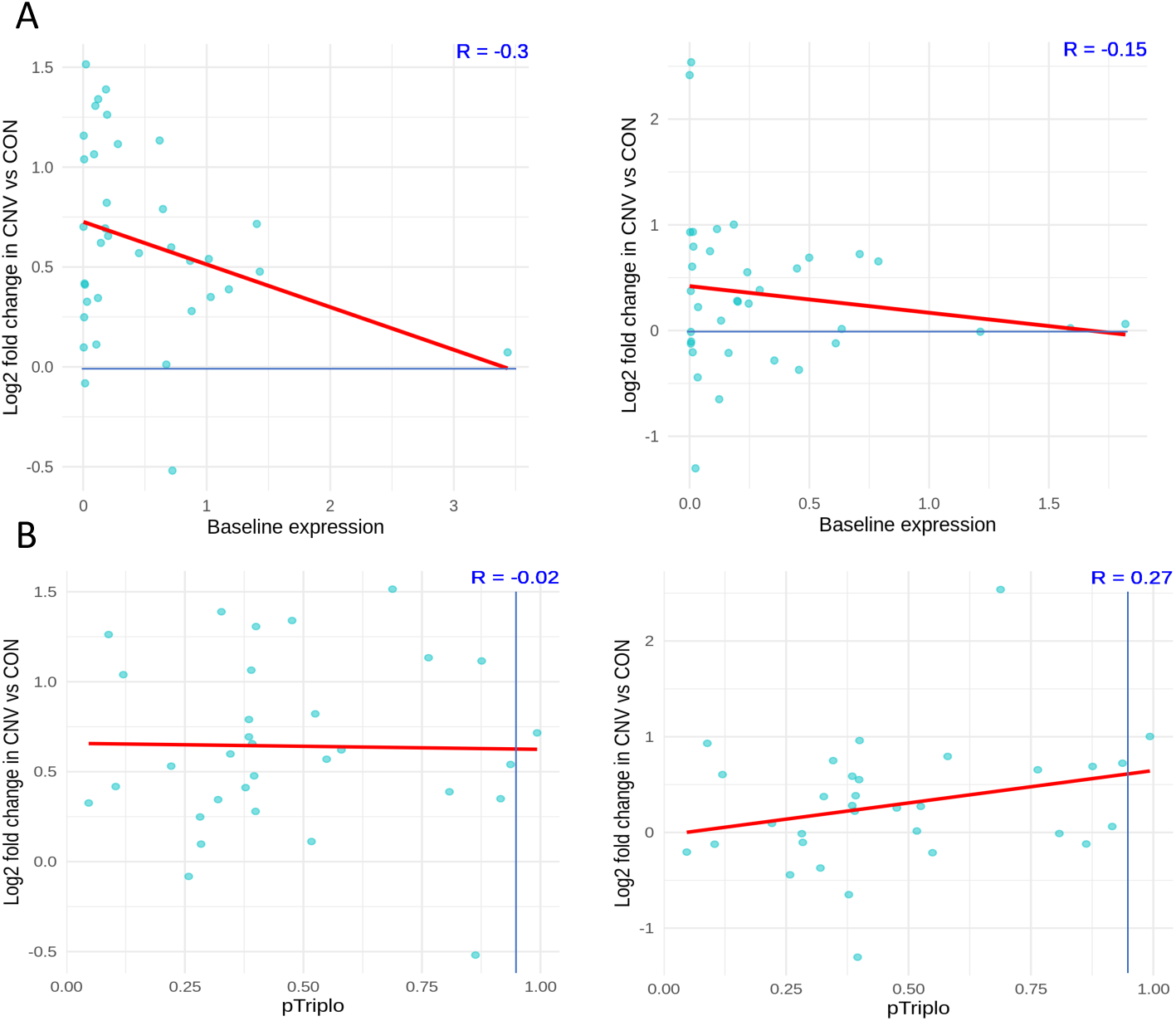
dup15q expression changes are not explained by baseline expression or genome-wide metrics. **A.** Relationship of baseline expression of dup15q genes and fold change in neurons (left) vs glia (right). Regardless of cell type, genes with high baseline expression show modest increases in dup15q cases. **B.** Relationship of pTriplo metric and fold change of dup15q genes in neurons (left) and glia (right). pTriplo > .94 is categorized as “triplosensitive”, note dearth of dup15q genes meeting this criteria.

Given that not all genes within a CNV are necessarily critical to the phenotype, we sought to clarify which genes would be both dysregulated and critical to the dup15q phenotype. To do this we made use of recent advances in quantifying genome-wide dosage sensitivity to overexpression. Specifically, the ‘pTriplo’ metric or the probability of triplosensitivity, was recently developed from a cross-disorder catalog of rare CNVs that confer susceptibility to human disease, where a number closer to one indicates a higher likelihood of being triplosensitive^34^. We hypothesized that many genes within the locus, particularly the PWACR, previously deemed critical for the clinical phenotype, would demonstrate a high triplosensitive score, and a subset of those furthermore would show highest fold change in expression in dup15q cases. Surprisingly, most genes of interest within the locus did not meet criteria as being triplosensitive as defined by the cutoff score of > .94. (Fig S5). Furthermore, genes with higher triplosensitivity metric again showed modest gene expression changes, suggestive of homeostatic regulatory mechanisms at play, regardless of cell type examined.

Carriers of dup15q have higher rates of epilepsy ^5,35^, and in fact all of the dup15q cases analyzed here have a history of this co-morbidity. To determine whether this might be influencing gene expression changes within the region of the duplication, we separated out the (non-dup15q) ASD cases by the presence or absence of seizure diagnosis. We observed no global difference between presence of an epilepsy/seizure diagnosis and absence in non-dup15 ASD within our region of interest, and neither of these sub-groups resembled dup15q cases, arguing against the presence of seizure as a being a major confounding factor (Fig S6).

Our dataset also affords an opportunity to compare analyses presented here to independent cell-type-specific reports on ASD. Encouragingly, we replicated past findings of neurodevelopmental disorders in our own gene expression data, including enrichment of known pathogenic ASD risk genes, FMR1 protein target genes, and dysregulation of layer 2/3 excitatory neurons (Fig S7A-B) ^7,9,27,30^. Using gene ontology analysis, we also found evidence for biological processes involved in synaptic function, in both excitatory and inhibitory subtypes in ASD and dup15q (Fig S7C).

We examined global changes in DNA accessibility using snRNA-seq and snATAC-seq samples (i.e. multi-omic sequencing) (Fig 5A), by identifying differentially accessible genomic regions between conditions (Fig 5B, Supplemental File 7-8) and overrepresented motifs (Fig 5C, Supplemental File 9-11). Although gene expression analysis showed a greater number of DEGs in dup15q versus control, as compared to non-dup15q ASD versus control, we identified twice as many differentially accessible *peaks* in non-dup15q ASD versus control brains (Fig 5B), consistent with dup15q primarily impacting a specific genetic region. Additionally, there were almost 4x as many differentially accessible peaks that were unique to either comparison, suggesting divergent patterns of genome-wide regulation in ASD and dup15q (Fig 5B).

**Figure 5.**
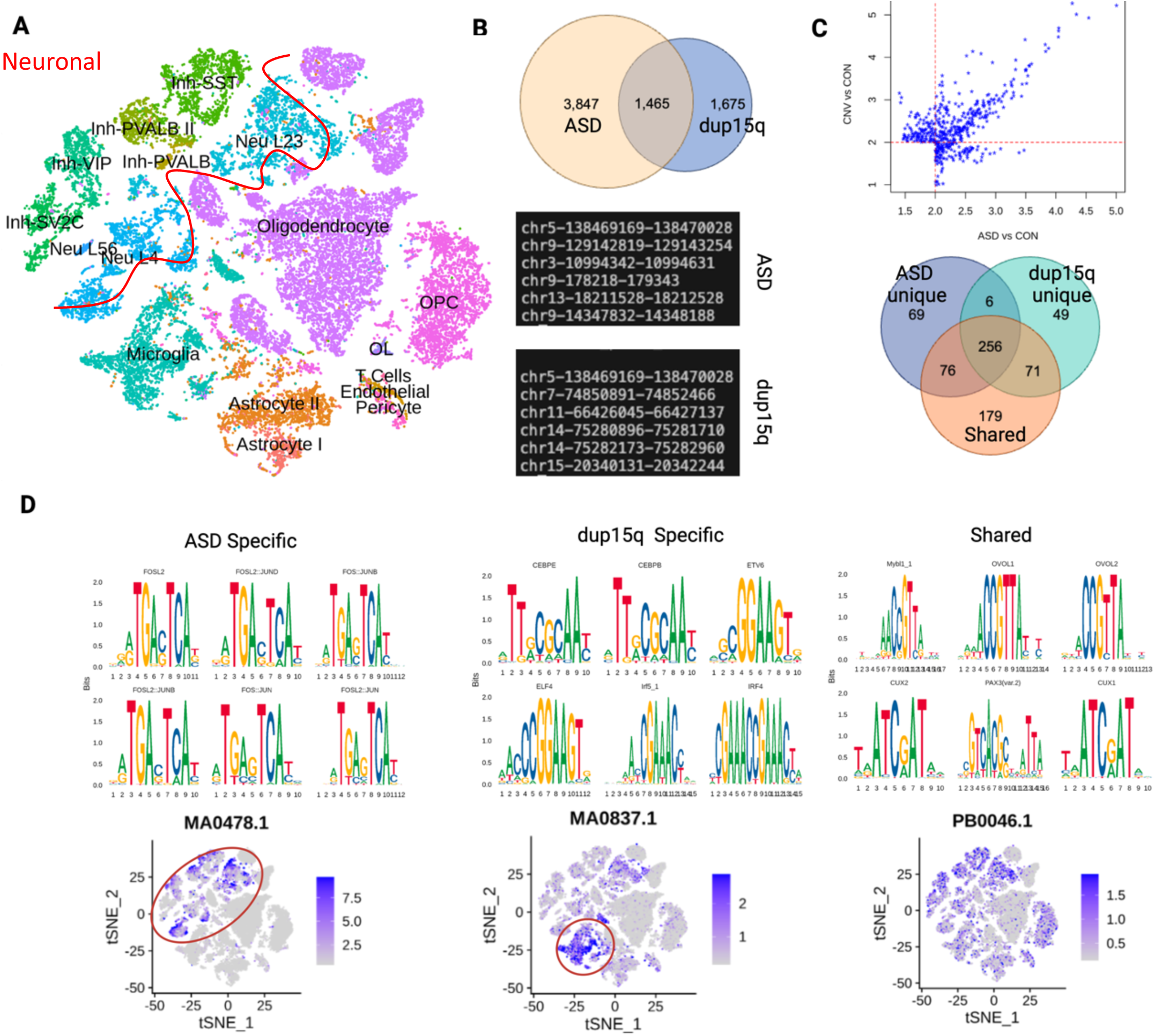
Multi-omic analysis reveals cell-type specific transcriptional regulatory activity in dup15q cases. **A.** Visualization of integration of snRNA-seq and snATAC-seq. **B.** Differential binding analysis revealed twice as many unique peaks in ASD vs control as compared to dup15q cases vs. control. Examples of genomic locations for ASD and dup15q shown, note that dup15q contains loci both within and outside of the duplicated region. **C.** The identified peaks in B were used to assess for fold enrichment of overrepresented motifs within differentially accessible genomic regions. Venn diagram demonstrates overlap of significant (i.e. fold enrichment >2 and padj <.05) of identified overrepresented motifs. **D.** FOS/JUN motifs were identified in the ASD specific differentially accessible genomic regions. Per cell motif activity calculation revealed higher activity in neuronal subtypes (circled). Conversely, motifs involved in immune and inflammation signaling were enriched in peaks unique to dup15q, with corresponding microglial activity (circled). Motifs enriched from the ‘shared’ group, (i.e. identified from differentially accessible sites present in both dup15q vs control and ASD vs control) lacked this cell-type-specificity. Representative JASPAR matrix profiles MA0478.1 (FOSL2), MA.0837.1 (CEBPE) and PB0046.1( Mybl1_1) shown.

Understanding the regulatory modules in ASD and dup15q is important to clarify the underlying neurobiology. For example, our analyses suggested that specific genomic regions demonstrating differential accessibility are largely unique in ASD and dup15q, there could be shared transcriptional regulatory modules due to convergent biological processes. We thus assessed enrichment for specific transcription factor motifs in the differentially accessible genomic regions that were unique to each condition vs those that were shared. The majority of motifs were not unique to any peak list (Fig 5C); these 256 motifs likely represent shared developmentally critical hubs of transcriptional regulation. This is consistent with their activity enrichment across different cell types in our data-set (Fig 5D). Among the top motifs that were unique to ASD (69 total) were FOS and JUN related sites; genome-wide analysis demonstrated these sites showed preferential activation in neuronal subtypes (Fig 5D). Examining the top enriched motifs unique to dup15q (49 total) demonstrated a distinct pattern-motifs critical to immune and inflammatory signaling were evident, such as CEBPE, ELF4 and IRF4. In contrast to the top ASD-unique motifs that demonstrated neuronal activity enrichment, our analyses highlighted microglial activity of these dup15q-unique motifs (Fig 5D). Furthermore, our snRNA-seq data showed that among non-neuronal subtypes, microglia contained the highest number of differentially expressed genes in dup15q (Supplemental Table 3). Additionally, gene ontology results for microglia in dup15q vs control revealed biological processes relevant both to inflammation as well as synaptic pruning, the latter of which was absent from the ASD vs control comparison (Fig_20_ S8). Thus, we uncovered an unexpected inflammatory signature unique to dup15q cases.

## Discussion

Here, we confirmed prior bulk RNA-seq work showing overall increased expression of genes within the region duplicated in dup15q brain ^36^ but further identified unexpected cell-type-specific heterogeneity in gene expression changes of genes within the duplicated region, specifically implicating dysregulation of the PWACR in neurons. Gene expression changes demonstrated different patterns of regulation at different genes not fully explained by gene dosage, function, or even imprinting status. Other reports of dup15q derived neurons demonstrated that cell type rather than genotype was the main determining factor in gene expression changes, findings with which our work agrees with ^30,37^. Unexpectedly, we found that the most highly expressed genes in any given cell-type seemed to be spared from the most profound effects of increased gene dosage and that prior genome-wide metrics of dosage sensitivity did not necessarily capture gene expression changes. Our work thus highlights the importance of unbiased and creative approaches in functionally validating the effects of copy number variants directly in the tissue of interest in neurodevelopmental disorders^38,39^.

We also identified transcriptional regulatory networks implicating neuroinflammation and microglial function. However, microglial *CYFIP1*, a gene within the duplicated region, did not demonstrate evidence for significant regulation in dup15q cases, despite our study being well-powered to detect such changes in this cell type. Our work, which is observational in nature, cannot parse whether the microglial signal observed here represents a direct effect of the duplication or a response by the brain to dysfunction in other cell types. Future experimental work will be required to address this. This finding is nonetheless intriguing in light of the emerging role of microglia in brain disorders across the lifespan^33,40–42^. We also resolved the cell-type-specific changes in expression of the GABA receptor cluster within the duplicated region. Given findings that individuals with dup15q have an EEG biomarker mimicking increased GABAergic signaling, as well as recent clinical trials which target GABA transmission, this information may provide useful in guiding future precision therapeutics^43,44^.

Reassuringly, our work is concordant with past studies, which reinforces the biological plausibility of our novel and unexpected findings of cell-type-specificity of dup15q. For example, our multi-omic sequencing revealed enrichment of JUN and FOS motifs in differentially accessible regions in ASD but not dup15q. This is consistent with an existing literature implicating these networks in neurodevelopmental disorders including ASD ^45,46^. Replication of independent findings of enrichment of SFARI ASD risk genes and FMR1 protein networks, as well as layer 2/3 neuronal dysregulation in the pathogenesis of neurodevelopmental disorders also indicated that our approach detected meaningful biological perturbations^7,9,30,36,27^. Indeed, results here confirmed work from independent laboratories using cellular models, as well as past experiments in human brain bulk tissue ^6,37^.

Our data show that the effects of copy number duplications are not straightforward. Given that dup15q is a germline event, it was unexpected that neurons demonstrated more marked dysregulation of gene expression within the duplicated region than glia. Although this implicates neuronal dysfunction in dup15q pathogenesis, it also provokes questions about the endogenous factors related to the epigenetic landscape that moderate the effects of this germline structural variant. It is interesting that we found that cell types with high gene expression at baseline may conversely engage homeostatic mechanisms to minimize further increases, which could be potentially toxic to the cell. Better understanding of these differences could be used in the future to guide therapeutic strategies, for example, by harnessing glial homeostatic mechanisms to tamp down effects of the duplication in neurons. Our unbiased, empiric identification of cell-type-specific changes in gene expression and the epigenetic landscape can also guide future therapeutic work. For example, non-canonical cell types may be critical in mediating the pathophysiology of dup15q, and thus benefit from targeting in future precision medicine therapeutic interventions, results that could easily be otherwise missed.

Limitations of this report include a small sample size of cases. Thus, certain features, such as individual variability of dup15q or the effects of the duplication on very rare cell types, cannot be completely parsed from our methodological approach. However, our main findings, of unexpected cell-type-specific determinants of the effect of dup15q, are unlikely to be artifacts of our sample size. Our ‘low-resolution’ analysis of glia and neurons corroborate the findings of the more subtle ‘high-resolution’ approach. It is possible that the unique microglial inflammatory signature in dup15q is an artifact of the relatively high nuclei number that this cell type comprises. This is unlikely given other cell types are present at similar or even higher proportions (astrocytes, for example), and were not implicated in multi-omic analysis in the same way. Future work will be needed to study whether the changes we identified are a direct result of alterations in the integrity of the imprinting center or more general genomic alterations. As one of the first reports of the cell-type-specific effects of copy number gains in human brain in neurodevelopmental disorders, our work reveals unexpected changes in gene expression that may fuel novel paths of investigation for therapeutic approaches. We also demonstrate the importance of direct, unbiased, and cell-type-specific studies in human brain to complement mouse and *in vitro* cellular models.

## Conclusions

We identified marked cell-type-specific regulation of gene expression and chromatin accessibility in human brain specimens from dup15q vs. non-dup15q ASD and neurotypical controls, both within the region of the duplication and genome-wide. This work demonstrates gene expression changes caused by copy number variants in neurodevelopmental disorders result from a complex interplay of factors as opposed to simply reflecting gene dosage. Neurons demonstrate marked evidence of altered gene expression in dup15q, particularly within the PWACR. We also identified evidence of an unexpected microglial inflammatory response unique to dup15q. This unexpected pattern of genomic regulation demonstrates the importance of directly studying functional effects of germline genetic structural variants in human brain in neurodevelopment, particularly in the context of copy number gains which may behave unexpectedly. Our findings also demonstrate the importance of studying primary brain tissue directly with techniques that afford cell-type-specificity and unbiased approaches. Our work will guide future mechanistic studies as well as rational therapeutic development in dup15q.

## List of abbreviations

ASD: autism spectrum disorder
CNV: copy number variant
PW/ACR: Prader-Willi/Angelman critical region
snRNAseq: single-nucleus RNA-sequencing
UMI: unique molecular index

## Acknowledgements

We are grateful and indebted to the families who donated tissue for research purposes to Autism BrainNet, the ATP and the NIH Neurobiobank at the University of Maryland, Baltimore, MD. The Autism BrainNet is a resource of the Simons Foundation Autism Research Initiative (SFARI). Autism BrainNet also manages the Autism Tissue Program (ATP) collection, previously funded by Autism Speaks. We thank Jennifer E. Neil for assistance with human samples; Robert Sean Hill, Dilenny Gonzalez, Shyam Akula and Sattar Khoshkoo for assistance in reagent ordering and sample sequencing; Sara Bizzoto and Sattar Khoskkoo for discussion of cell type specific markers in cortex, Ronald Mathieu and the Hematology/Oncology Flow Cytometry Research Facility for assistance with sorting; the Engle lab and the Harvard Biopolymers Facility for assistance with Chromium Controller use and sequencing; Emre Caglayan, Martin Breuss and Tamim Shaikh for thoughtful comments on the manuscript. Molecular genetics library quantification support was provided by the Boston Children’s Hospital Intellectual and Developmental Disabilities Research Center Molecular Genetics Core Facility supported by U54HD090255 from the NIH Eunice Kennedy Shriver National Institute of Child Health and Human Development. The bioinformatics analysis was performed with the computational resources provided by the Research Computing Group at Boston Children’s Hospital and Harvard Medical School, including High-Performance Computing Clusters Enkefalos 2 (E2), and the BioGrids scientific software made available for data analysis. This work was supported by a grant from the Simons Foundation and Autism BrainNet (953759, C.A.W). C.A.W. is also supported by the NIMH (U01MH106883) and the Tan Yang Center for Autism Research at Harvard Medical School. C.A.W. is an Investigator of the Howard Hughes Medical Institute. C.M.D. is a Boettcher Webb-Waring Early Career Investigator and was previously supported in part by NIMH Translational Post-doctoral Training Program in Neurodevelopment T32MH112510.

## Contributions

C.M.D. performed the snRNA-seq experiments, analyzed the data, and wrote the manuscript draft. A.M. conducted multi-omic sequencing experiments and analyzed the data. C.C., L.S., and S.W conducted bioinformatic analysis of transcriptomic and multi-omic sequencing. C.B. and K.C. conducted optical mapping experiments and analysis. C.A.W, C.M.D., and A.M. conceptualized the project, and C.A.W supervised the project. All authors reviewed and contributed to revising and editing the manuscript draft.

## SUPPLEMENTAL FIGURES

**Figure S1:**
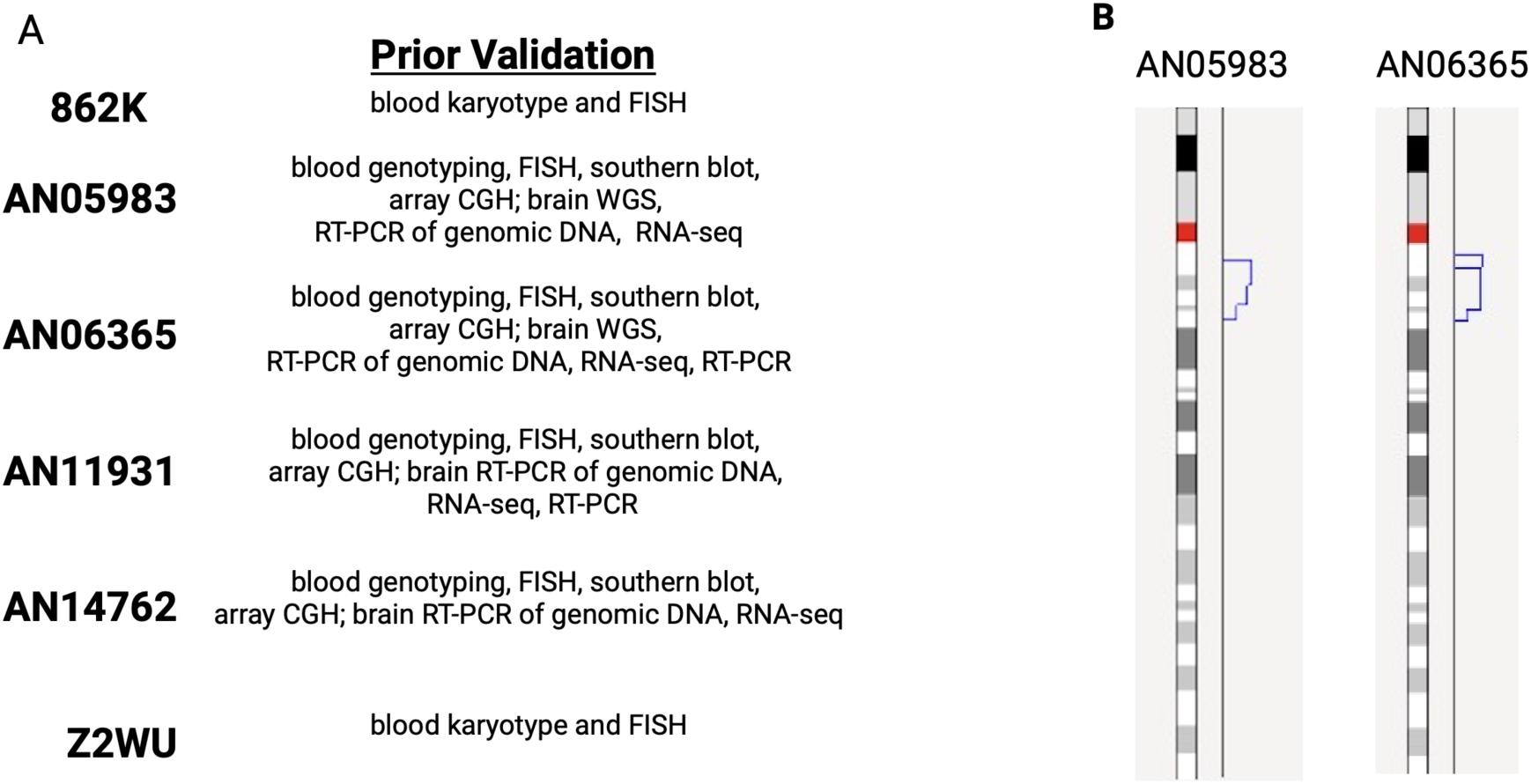
Dup15q sample validation (all samples have tetrasomy of the PWACR) **A.** Dup15q samples have been extensively validated in the literature^6,36^. **B.** Two samples were also validated with optical genome mapping. Note that the ideogram represents only the copy number changes and not the chromosomal structure.

**Figure S2:**
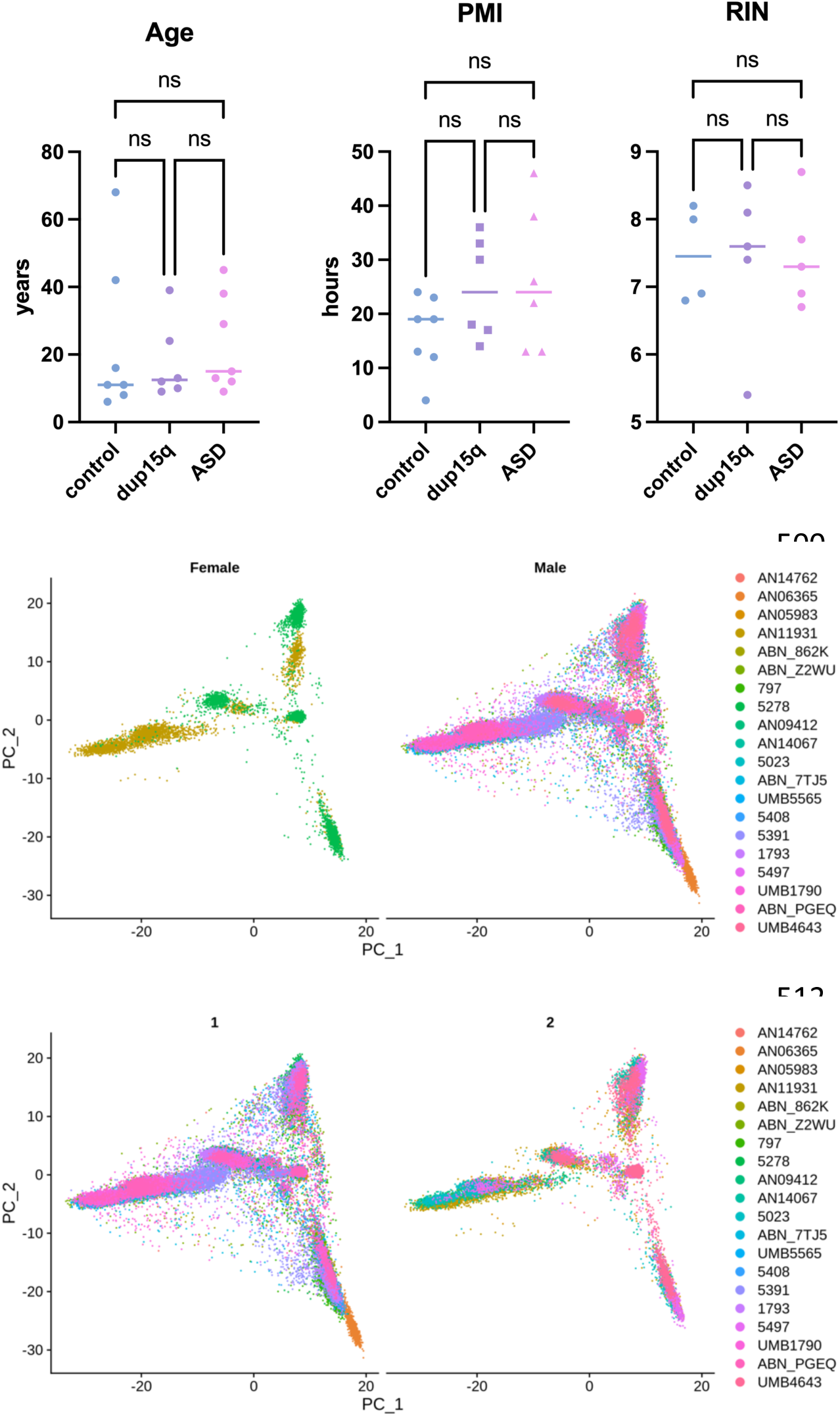
Top panel: Sample comparison. No significant differences in age, PMI or RIN. (p> .05, 1 way ANOVA, Tukey post-hoc test. There was also no difference in the female percentage between groups (p>.05, chi-square). Middle and bottom panel: PCA plots demonstrate sex and age (split as< and > 21 years of age) are not major determinants of gene expression variability.

**Figure S3:**
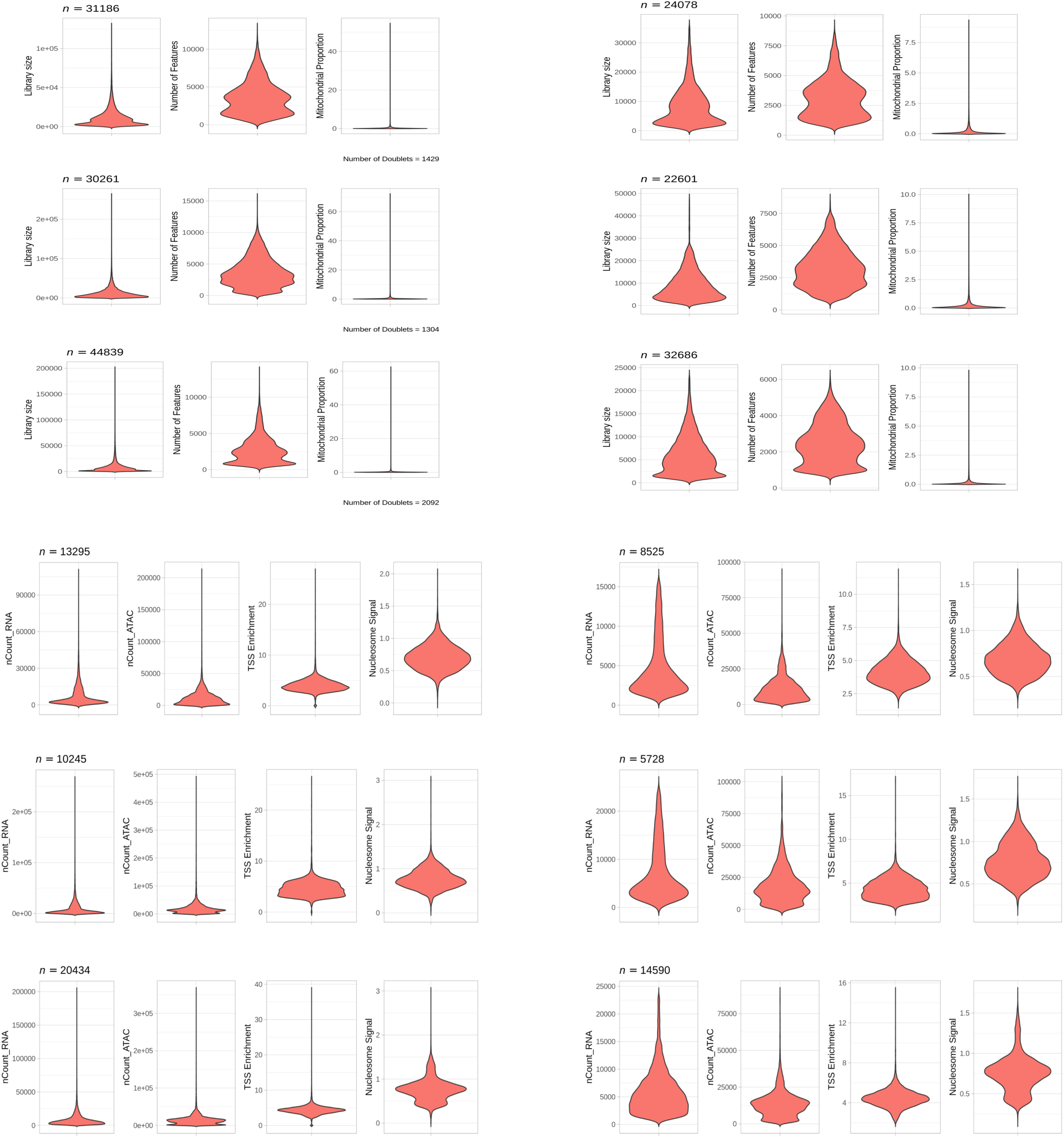
snRNA-seq (top 3) and multi-omic (bottom 3) quality metrics of raw (left) and filtered (right) nuclei. Each panel demonstrates library size, gene number, and proportion of mitochondrial genes (from left to right) and for ATAC-seq-ncount RNA, ATAC, TSS enrichment and nucleosome signal are shown. Top is dup15q, middle is ASD, and bottom is control. Number of nuclei in top left corner, number of doublets in each group prior to filtering also indicated.

**Figure S4:**
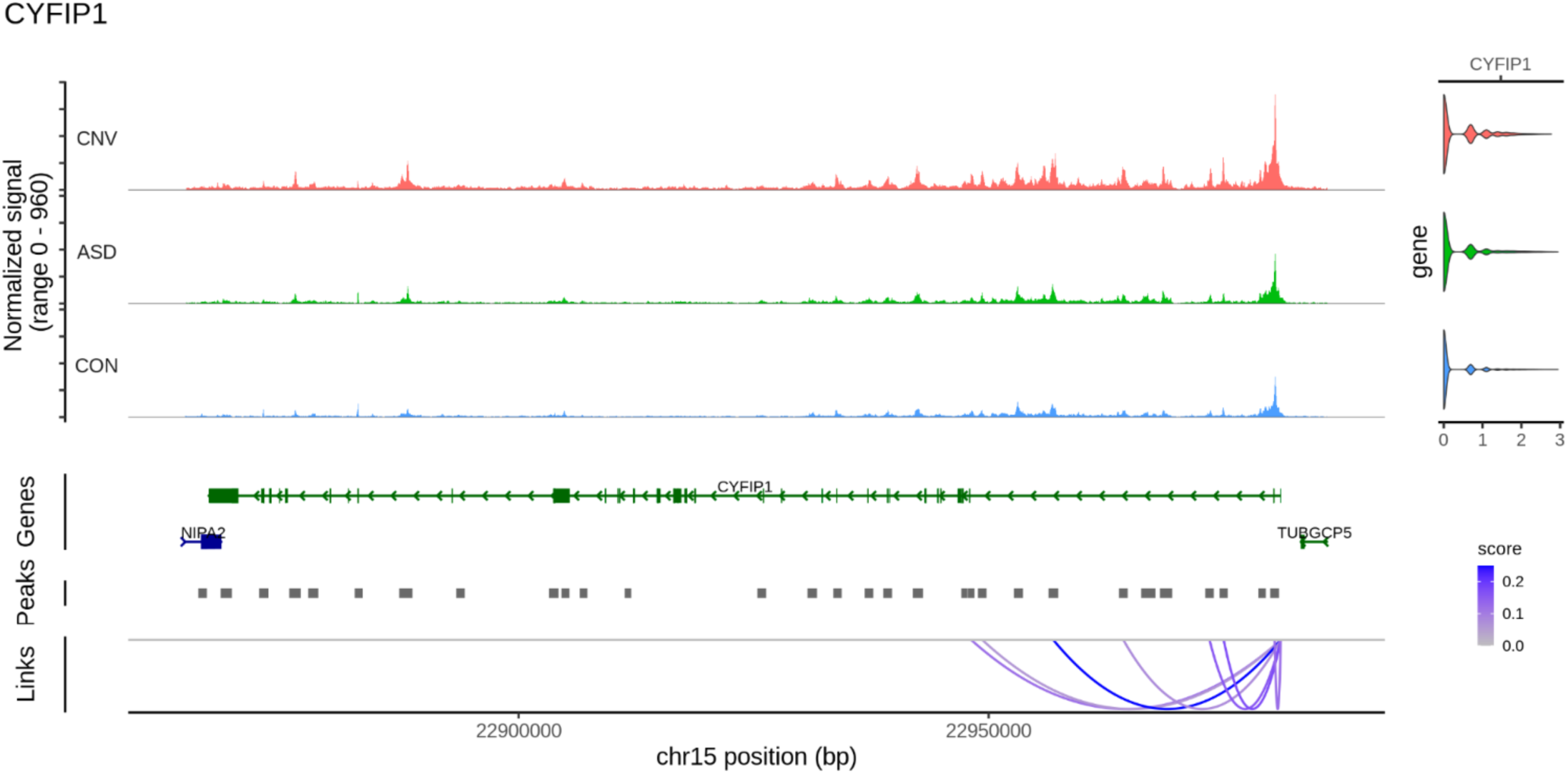
Increased expression and chromatin accessibility at *CYFIP1* in dup15q samples shown, as well as peak to peak linkage. Represents signal from all nuclei combined.

**Figure S5.**
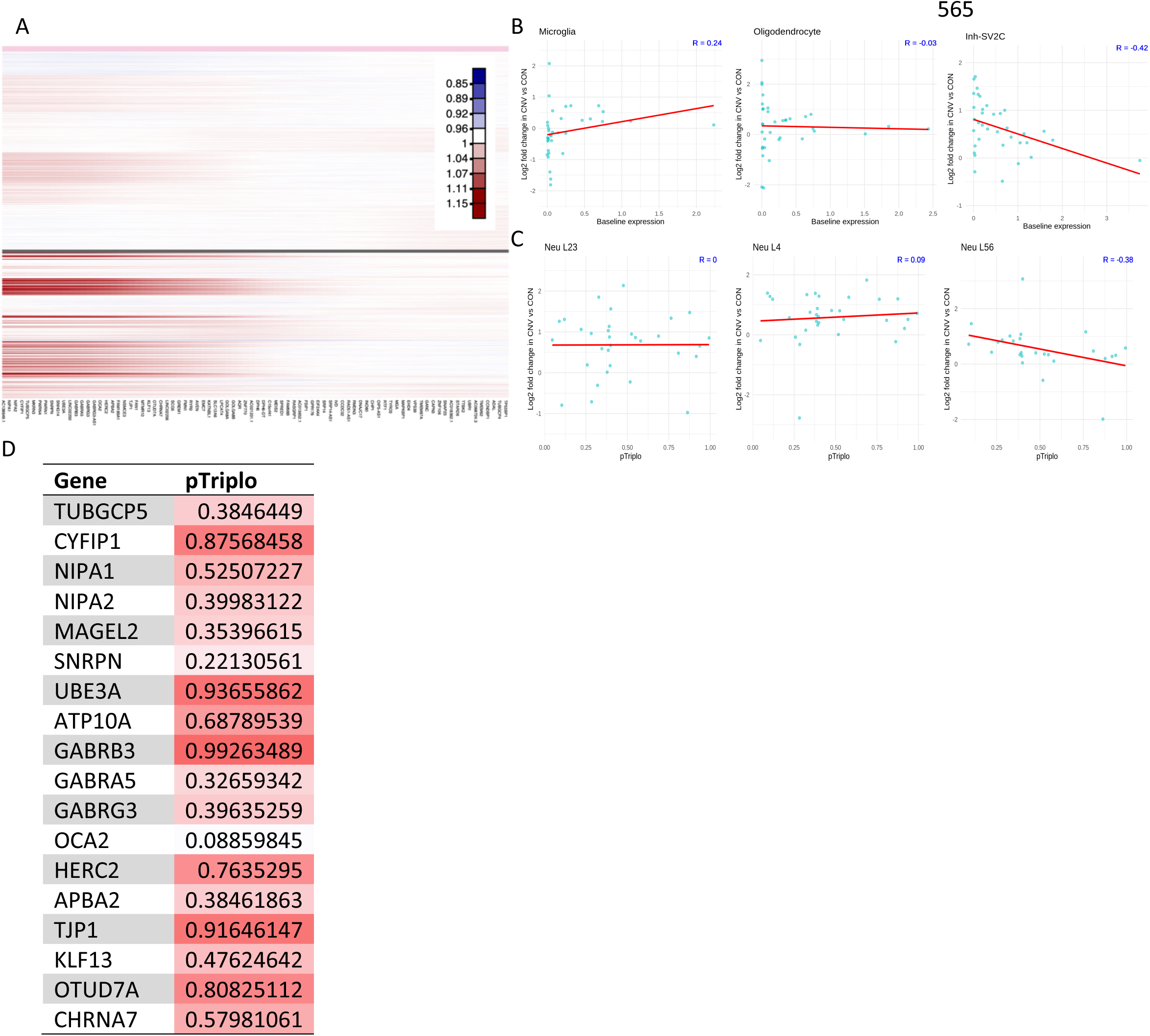
Assessing mediating factors to dup15q expression changes. **A.** inferCNV reveals heterogeneous expression increases in region of duplication in dup15q cases (below grey line) across samples as compared to control (above grey line) within all nuclei (each row). Heatmap color scale indicates decreased (blue) or increased expression. **B.** Cell-type-specific examples of the association between baseline expression in controls and the fold change expression in dup15q. Within the duplicated region, genes that are highly expressed in different cell types demonstrate modest changes in expression in dup15q cases. p-values are non-significant except for in Inh-SV2C (p=.01, not adjusted for multiple comparisons) **C.** Association of pTriplo in cell-type-specific examples. All p-values > .050. **D.** pTriplo metric for select dup15q genes.

**Figure S6:**
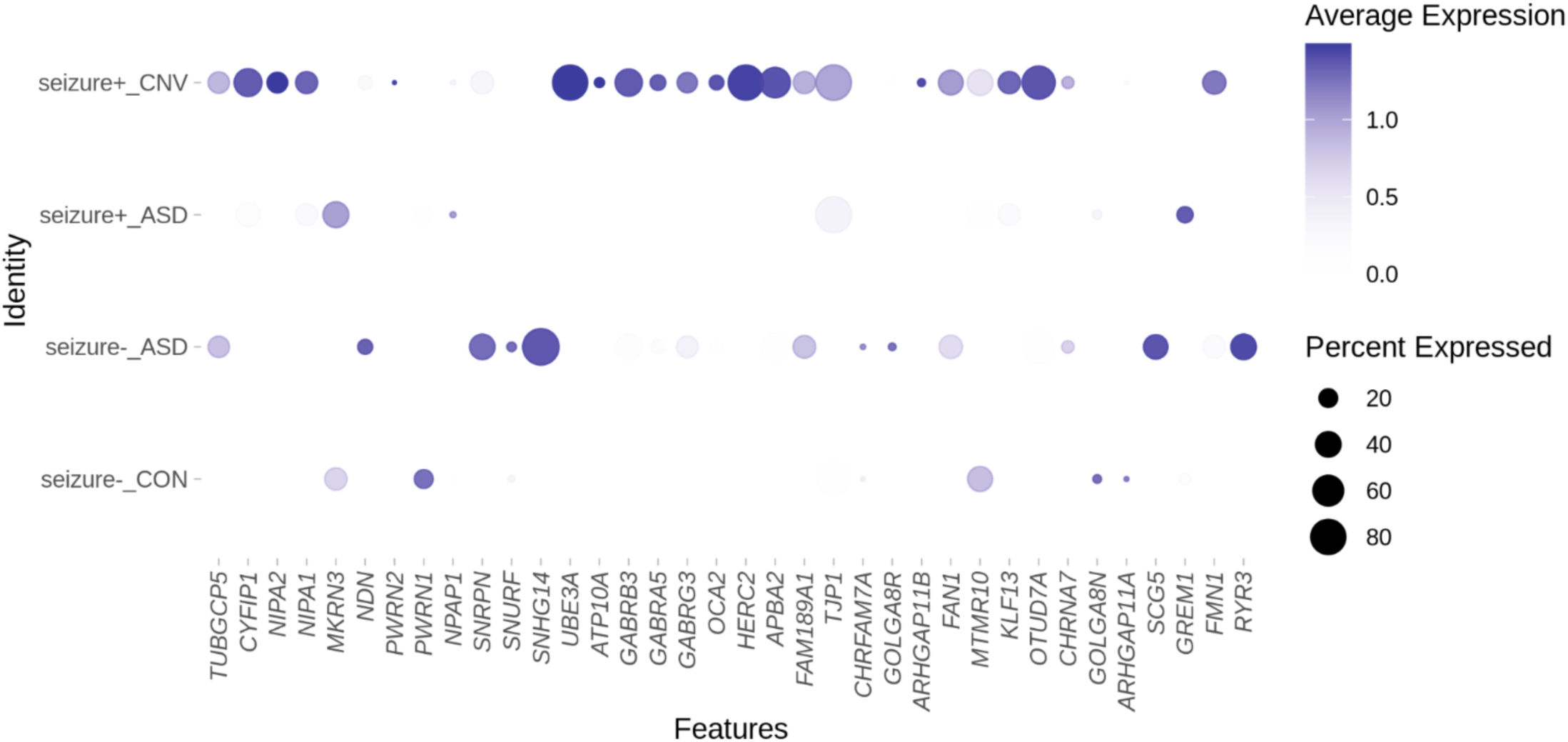
Presence of seizure diagnosis in ASD cases does not recapitulate dup15q gene expression changes observed within the duplicated (CNV) region.

**Figure S7:**
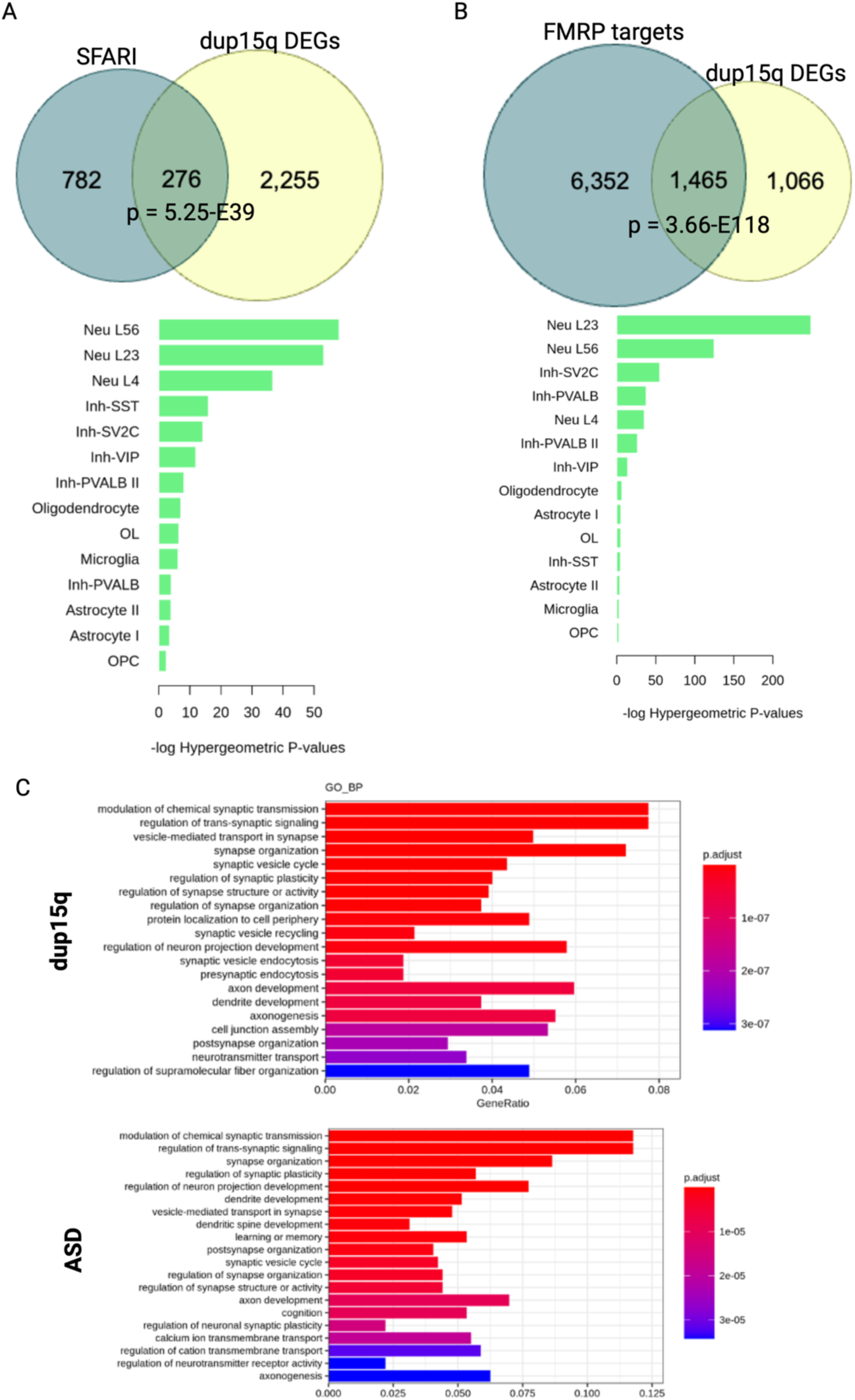
Overlap of known (A) SFARI pathogenic ASD risk genes as well as (B) FMR1 protein target genes, demonstrates significant enrichment (p-value indicated in figure, hypergeometric test) These genes demonstrate notable enrichment in excitatory neuron subtypes. **C.** Using gene ontology analysis, we also found evidence for biological process terms involved in synaptic function. Presented are results from differential expression analysis in layer 2/3 neurons in each condition vs. control.

**Figure S8:**
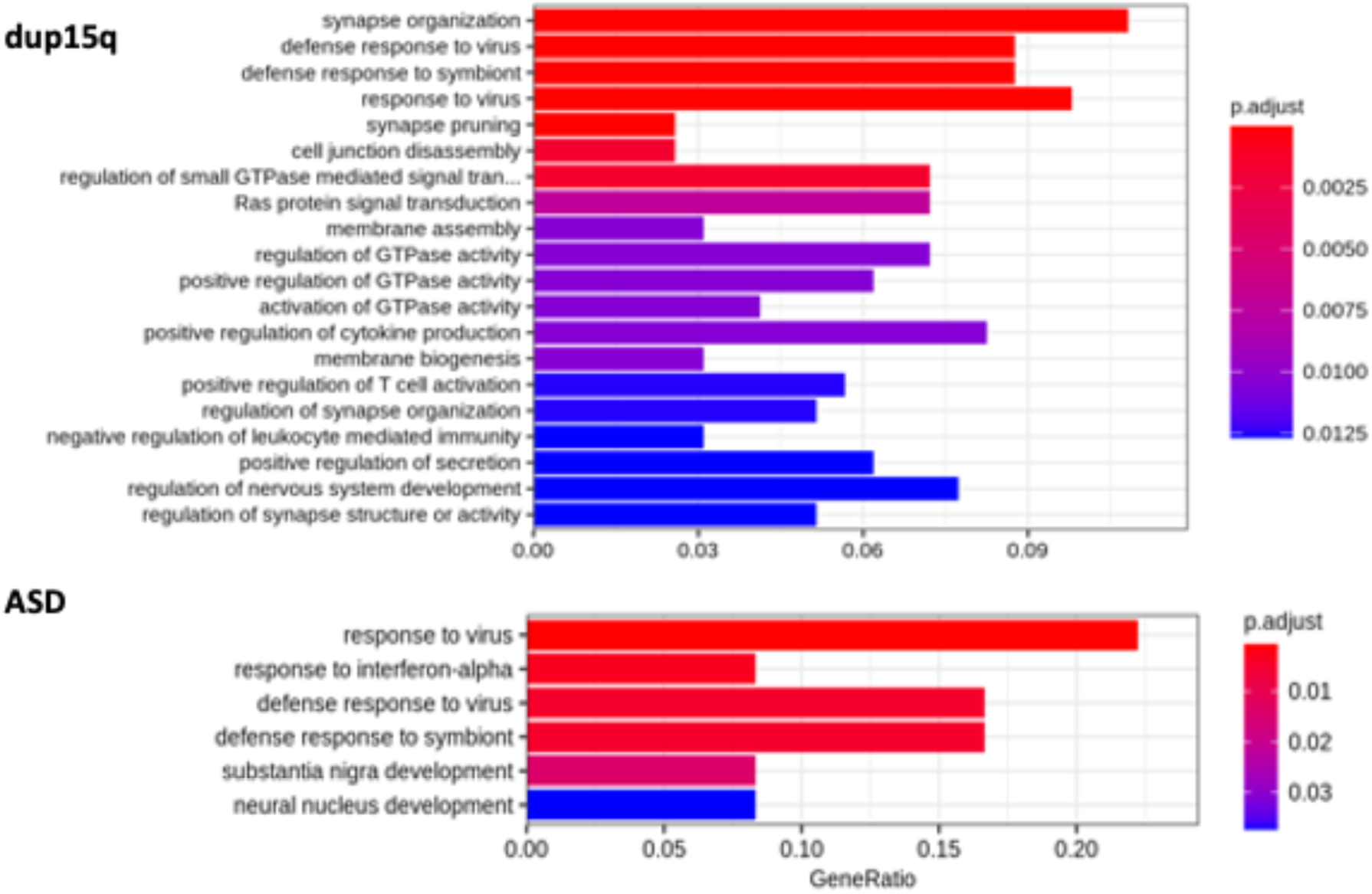
Using gene ontology analysis, we also found evidence for biological process terms related to inflammation and synaptic pruning in dup15q microglia, but not ASD microglia (compared to control)

**Supplemental Table 1:** Number of significant differentially expressed genes with different log fold change cut-off in global differential expression analysis (i.e. using all nuclei from different cell types combined).

| All Cells | # DEGs<br>(logfc=0.25) | # DEGs<br>(logfc=0.2) |
| --- | --- | --- |
| CNV vs CON | 47 | 126 |
| ASD vs CON | 6 | 34 |
| CNV vs ASD | 23 | 52 |

**Supplemental Table 2:** Number of significant differentially expressed genes with different log fold change cut-off in differential expression analysis separating neuron and glia.

| Neuron Cells | # DEGs<br>(logfc=0.25) | # DEGs<br>(logfc=0.2) |
| --- | --- | --- |
| CNV vs CON | 259 | 623 |
| ASD vs CON | 111 | 329 |
| CNV vs ASD | 38 | 101 |

| Glia Cells | # DEGs<br>(logfc=0.25) | # DEGs<br>(logfc=0.2) |
| --- | --- | --- |
| CNV vs CON | 95 | 206 |
| ASD vs CON | 17 | 45 |
| CNV vs ASD | 35 | 113 |

**Supplemental Table 3:** Number of significant differentially expressed genes in high resolution differential expression analysis, in dup15q vs control comparison (p-value adj<0.05).

| Cell Type | # DEGs | # Upreg | # Downreg | Cell Type | # DEGs | # Upreg | # Downreg |
| --- | --- | --- | --- | --- | --- | --- | --- |
| Non-neuronal |  |  |  | Neuronal |  |  |  |
| Pericyte | 48 | 23 | 25 | Neu L56 | 3 | 2 | 1 |
| Endothelial | 11 | 7 | 4 | Neu L4 | 308 | 239 | 69 |
| Microglia | 240 | 124 | 116 | Neu L23 | 1,307 | 1,162 | 145 |
| Astrocyte I | 115 | 60 | 55 | Inh-SV2C | 315 | 290 | 25 |
| Astrocyte II | 124 | 82 | 42 | Inh-VIP | 107 | 84 | 23 |
| Oligodendrocyte | 119 | 78 | 41 | Inh-PVALB | 175 | 165 | 10 |
| OL | 5 | 3 | 2 | Inh-SST | 74 | 40 | 34 |
| OPC | 22 | 14 | 8 | Inh-PVALB II | 205 | 193 | 12 |
| T Cells | 1 | 1 | 0 |  |  |  |  |

